# The tumor microbiome as a predictor of outcomes in patients with metastatic melanoma treated with immune checkpoint inhibitors

**DOI:** 10.1101/2023.05.24.542123

**Authors:** Caroline E. Wheeler, Samuel S. Coleman, Rebecca Hoyd, Louis Denko, Carlos H.F. Chan, Michelle L. Churchman, Nicholas Denko, Rebecca D. Dodd, Islam Eljilany, Sheetal Hardikar, Marium Husain, Alexandra P. Ikeguchi, Ning Jin, Qin Ma, Martin D. McCarter, Afaf E.G. Osman, Lary A. Robinson, Eric A. Singer, Gabriel Tinoco, Cornelia M. Ulrich, Yousef Zakharia, Daniel Spakowicz, Ahmad A. Tarhini, Aik Choon Tan

## Abstract

Emerging evidence supports the important role of the tumor microbiome in oncogenesis, cancer immune phenotype, cancer progression, and treatment outcomes in many malignancies. In this study, we investigated the metastatic melanoma tumor microbiome and potential roles in association with clinical outcomes, such as survival, in patients with metastatic disease treated with immune checkpoint inhibitors (ICIs). Baseline tumor samples were collected from 71 patients with metastatic melanoma before treatment with ICIs. Bulk RNA-seq was conducted on the formalin-fixed paraffin-embedded (FFPE) tumor samples. Durable clinical benefit (primary clinical endpoint) following ICIs was defined as overall survival ≥24 months and no change to the primary drug regimen (responders). We processed RNA-seq reads to carefully identify exogenous sequences using the {exotic}tool. The 71 patients with metastatic melanoma ranged in age from 24 to 83 years, 59% were male, and 55% survived >24 months following the initiation of ICI treatment. Exogenous taxa were identified in the tumor RNA-seq, including bacteria, fungi, and viruses. We found differences in gene expression and microbe abundances in immunotherapy responsive versus non-responsive tumors. Responders showed significant enrichment of several microbes including *Fusobacterium nucleatum*, and non-responders showed enrichment of fungi, as well as several bacteria. These microbes correlated with immune-related gene expression signatures. Finally, we found that models for predicting prolonged survival with immunotherapy using both microbe abundances and gene expression outperformed models using either dataset alone. Our findings warrant further investigation and potentially support therapeutic strategies to modify the tumor microbiome in order to improve treatment outcomes with ICIs.

**Significance:** We analyzed the tumor microbiome and interactions with genes and pathways in metastatic melanoma treated with immunotherapy, and identified several microbes associated with immunotherapy response and immune-related gene expression signatures. Machine learning models that combined microbe abundances and gene expression outperformed models using either dataset alone in predicting immunotherapy responses.

## INTRODUCTION

Advances in immunotherapy, including immune checkpoint inhibitors (ICIs), have transformed the standard of care for many types of cancer, including melanoma. While ICIs have improved outcomes for melanoma patients, many patients suffer from primary or secondary tumor resistance. For example, at 6.5 years, the overall survival rates with ipilimumab plus nivolumab, nivolumab, and ipilimumab were 49%, 42%, and 23%, respectively, as reported in the pivotal CheckMate 067 trial (1). Furthermore, mechanisms of resistance to immunotherapy remain poorly understood, and many treatments are associated with immune-mediated toxicities. Therefore, there is an urgent need to develop and improve biomarkers predictive of benefit from ICI therapy.

Numerous biomarkers that predict the response of melanoma to ICIs are under investigation, including those based on clinical characteristics, genomics, transcriptomics, and epigenomics. For genomics data, these predictive biomarkers include tumor mutational burden (TMB) (2), neoantigen load (3), genotypes of HLA-I (3,4), T-cell repertoire (5), aneuploidy (also known as somatic copy number alterations, SCNAs) (6), and germline variations (7). On the other hand, predictive biomarkers derived from transcriptomics data include tumor oncogene expression signatures such as genes related to MYC (8), WNT/ß-catenin (9,10), or RAS (11) signaling, or gene expression profiles within the tumor immune microenvironment (TIME) such as interferon-γ (IFN-γ) responsive genes (12), chemokines (13,14), major histocompatibility complex (MHC) class I and II (15), and cytotoxic T-cell and T-cell effector (16,17) gene expression markers that have been reported to be predictive of ICI response in metastatic melanoma. Unfortunately, the predictive power remains low. For example, in terms of prediction of ICI response, TMB, IFN-γ-responsive gene signatures, or the combination of TMB and IFN-γ gene signatures produce an area under the receiver operating characteristic curve (AUROC) of 0.60-0.84 in melanoma cohorts (18).

Recently, high-throughput transcriptome, genome, or amplicon-based sequencing data demonstrated an abundance and variety of microbes’ nucleic acids inside tumors (8). In some cases, hundreds of negative controls and paraffin-only blocks were sequenced to ensure a thorough understanding of the background signal and reagent contamination. Further, the presence of microbes has been validated using fluorescence in situ hybridization (FISH) and immunohistochemistry (IHC) (19). The microbes showed cancer specificity (9,12,13), and blood-based measurements could predict early-stage disease. These findings suggest that microbes observed in high-throughput sequencing data may also correlate with treatment outcomes.

Recent efforts to use these microbes as biomarkers showed that while generally less predictive of prognosis than gene expression, when combined with gene expression they increase the predictive power (20). Further, the tumor microbiome was predictive of chemotherapy response.

Here, we describe the use of tumor RNA sequencing (RNA-seq) to predict response to ICIs in patients with melanoma (**Figure 1**). We demonstrate the presence of microbes within tumors and show the existence of different microbial communities in patients whose tumors responded to treatment. We predict treatment response using human gene expression patterns that perform similarly to other ICI-response prediction efforts. Finally, we show how the presence of microbes correlates with these signatures, suggesting an interaction with the immune system, and how including tumor microbes in these models improves their predictive accuracy.

**Figure 1.**
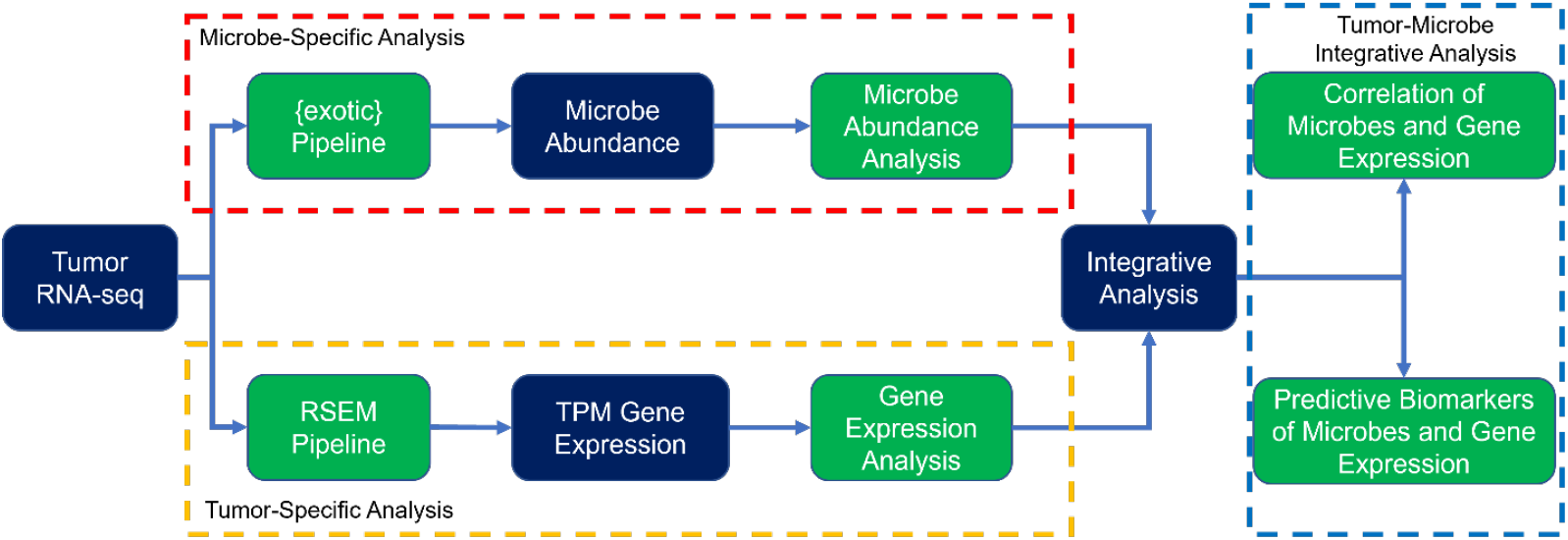
Overview of the research strategy. RNA-seq data from tumor specimens are processed to microbe abundances and human gene expression. Each is associated with IO response, and then integrative analyses combine them into a model to predict outcomes.

## MATERIALS AND METHODS

### Study design

Established in 2014, the Oncology Research Information Exchange Network (ORIEN) is an alliance of 18 US cancer centers. All ORIEN alliance members utilize a standard IRB-approved protocol: Total Cancer Care® (TCC). As part of the TCC, participants agree to have their clinical data followed over time, to undergo germline and tumor sequencing, and to be contacted in the future by their provider if an appropriate clinical trial or other study becomes available (21). TCC is a prospective cohort study where a subset of patients elect to be enrolled in the ORIEN Avatar program, which provides research use only (RUO)-grade whole-exome tumor sequencing, RNA-seq, germline sequencing, and collection of deep longitudinal clinical data with lifetime follow-up. Nationally, over 325,000 participants have enrolled in TCC. M2GEN, the commercial and operational partner of ORIEN, harmonizes all abstracted clinical data elements and molecular sequencing files into a standardized, structured format to enable the aggregation of de-identified data for sharing across the network. Data access was approved by the IRB in an Honest Broker protocol (2015H0185) and Total Cancer Care protocol (2013H0199) in coordination with M2GEN and participating ORIEN members.

In this study, we assembled RNA-seq data from the tumor samples of 71 patients with metastatic melanoma treated with ICIs. We defined durable clinical benefit (primary clinical endpoint) following ICIs as overall survival ≥24 months and no change to the primary drug regimen (hereafter referred to as responders).

### Sequencing methods

ORIEN Avatar specimens undergo nucleic acid extraction and sequencing at HudsonAlpha (Huntsville, AL) or Fulgent Genetics (Temple City, CA). For frozen and OCT tissue DNA extraction, Qiagen QIASymphony DNA purification is performed, generating a 213 bp average insert size. For frozen and OCT tissue RNA extraction, Qiagen RNAeasy plus mini kit is performed, generating 216 bp average insert size. For formalin-fixed paraffin-embedded (FFPE) tissue, a Covaris Ultrasonication FFPE DNA/RNA kit is utilized to extract DNA and RNA, generating a 165 bp average insert size. RNA-seq is performed using the Illumina TruSeq RNA Exome with single library hybridization, cDNA synthesis, library preparation, and sequencing (100 bp paired reads at Hudson Alpha, 150 bp paired reads at Fulgent) to a coverage of 100M total reads/50M paired reads.

### Data processing and gene expression analyses

RNA-seq Tumor Pipeline Analysis is processed according to the workflow outlined below using GRCh38/hg38 human genome reference sequencing and GenCode build version 32. Adapter sequences are trimmed from the raw tumor sequencing FASTQ file. Adapter trimming via k-mer matching is performed along with quality trimming and filtering, contaminant filtering, sequence masking, GC filtering, length filtering, and entropy filtering. The trimmed FASTQ file is used as input to the read alignment process. The tumor adapter-trimmed FASTQ file is aligned to the human genome reference (GRCh38/hg38) and the Gencode genome annotation v32 using the STAR aligner. The STAR aligner generates multiple output files for Gene Fusion Prediction and Gene Expression Analysis. RNA expression values are calculated and reported using estimated mapped reads, fragments per kilobase of transcript per million (FPKM) mapped reads, and transcripts per million (TPM) mapped reads at both the transcript and gene levels based on transcriptome alignment generated by STAR. RSEM pipeline and gene expressions were quantified as TPM. Gene expressions (GE) were log2(TPM+1) transformed, and downstream analyses were performed using the GE matrix. To determine differentially expressed genes (DEG) of responders vs. non-responders, we used the *limma* (v. 3.54.0) and *edgeR* (v. 3.40.0) packages where genes that have log2 fold change (log2FC) greater or less than 1 and adjusted p-value ≤0.1 were considered as significant DEG. For gene set enrichment analysis (GSEA) of responders vs. non-responders, we used the Java version of *gsea* (v. 4.3.2) using the gene set permutation of 1000 using Hallmark gene sets or TIMEx cell types. Gene sets or cell types that have adjusted p-value <0.1 were considered significant. Normalized enrichment score (NES) and adjusted p-value were provided in the plot.

### Microbe abundance and diversity

RNA-seq reads are used to calculate microbe abundances using the {exotic}pipeline, as described previously (22). Briefly, reads are aligned first to the human reference genome, and then unaligned reads are mapped to a database of bacteria, fungi, archaea, viruses, and eukaryotic parasites. The observed microbes then proceed through a series of filtering steps to carefully and conservatively remove contaminants before batch correction and normalization. Diversity measures were estimated by calculating the Shannon and Simpson indices, as well as Chao1, ACE, and inverse Simpson using the R package *vegan*.

### Signatures and pathways analyses

Gene signature scores were calculated using the IOSig and tmesig R packages. In brief, for each published gene signature, we collected and harmonized gene names using the NCBI Entrez gene number. To quantify the published gene expression score, we first transformed the gene expressions across samples within a cohort into a Z-score. Next, we averaged the standardized Z-score across the number of genes in the signature as previously described (15,23,24). This score is used to compare responders and non-responders of immunotherapies within individual cohorts based on the AUROC as previously described (23). We performed clustering of gene signatures based on the correlation of AUROC across multiple cohorts.

Within a cohort of patients, we stratified the patients into “high” or “low” groups based on the mean of the Z-score. A Mann-Whitney U test was performed in comparing the two groups to determine the difference, and the false discovery rate (FDR) of <0.05 was deemed to be significant. The list of published gene signatures are available as **Supplementary Table S1**.

For pathway analysis, single-sample GSEA (ssGSEA), via the ssGSEA method in the GSVA R package, was utilized to investigate the enriched gene sets in each sample. GSVA was run using the log2(TPM+1) gene expression values with Gaussian kernel. The Hallmark gene sets, TIMEx cell types, and the collected previously published gene expression signatures were used as the gene sets. The Hallmark gene sets are a curated list of gene sets that signify well-understood pathways that display reliable gene expression (25). The TIMEx cell types are formed from pan-cancer single-cell RNA-seq signatures and focus on illuminating immune cell infiltration from bulk RNA-seq data (26). A spearman correlation analysis was conducted using the differentially expressed microbe data and the 3 ssGSEA results. The gene sets were clustered according to the Euclidean distance with complete linkage, while the microbes were ordered from highest to lowest effect size.

### Prediction of response to treatment outcomes

To assess the predictive ability of the RNA-seq and microbe data for tumor response to ICIs, random forest classifiers were created using the *randomForest* R package. Models were based on 5 sets of input data: (1) microbe data, (2) 31-gene signature Z-score, (3) immune-activated gene signature Z-score, (4) microbe and 31-gene signature Z-score combined, and (5) microbe and immune-activated gene signature Z-score combined. Models were constructed with 500 trees and fivefold cross-validation. Additionally, 5 seeds were used for each model resulting in 25 trained models based on each set of input features. The AUROC curve was used to assess the overall performance of the trained models. This metric assesses the model classification accuracy, where 1 is a perfect classifier and 0.5 is a random classifier. The overall performance for each input feature-based model was taken as the average of the 25 trained models.

## RESULTS

### Patient Characteristics

From the ORIEN networks, we included 71 patients with metastatic melanoma in this study (IO_NOVA_Mel). The age of the patients in this cohort ranges from 24 to 83; 59% were male; and 55% survived >24 months following the initiation of ICI treatment (**Table 1**). ICI treatments included nivolumab (34.4% of non-responders, 10.3% of responders), pembrolizumab (25% of non-responders, 46.2% of responders), and others. Mean progression-free survival of responders (49.58 months) and non-responders (10.82 months) was significantly different (p-value <0.001).

**Table 1.** **Patient Demographics**, stratified by response to ICIs.

|  | NON-RESPONDER (N = 32) | RESPONDER (N = 39) | P-VALUE |
| --- | --- | --- | --- |
| AGE (MEAN (SD)) | 57.48 (15.85) | 58.62 (13.93) | 0.748 |
| SEX = MALE (%) | 18 (56.2) | 24 (61.5) | 0.835 |
| IO (N (%)) | **Atezolizumab** 1 (3.1)<br>**Ipilimumab** 6 (18.8)<br>**Ipilimumab + Nivolumab** 6 (18.8)<br>**Nivolumab** 11 (34.4)<br>**Pembrolizumab** 8 (25.0) | **Atezolizumab** 0 (0.0)<br>**Ipilimumab** 16 (41.0)<br>**Ipilimumab + Nivolumab** 1 (2.6)<br>**Nivolumab** 4 (10.3)<br>**Pembrolizumab** 18 (46.2) | 0.003 |
| RACE = WHITE (%) | 32 (100.0) | 39 (100.0) |  |
| PFS (MEAN (SD)) MONTHS | 10.82 (6.23) | 49.58 (19.24) | <0.001 |

### Gene expression analysis and its association with response to ICIs

Gene expression profiles for the 71 patients with metastatic melanoma treated with ICIs were obtained from ORIEN. We performed DEG analysis and identified five 5 genes (*CLEC12A, GBP1P1, CD96, CCL4, IDO1*) that were over-expressed in the responders as compared to the non-responders with log2FC >1 and adjusted p-value <0.1 (**Figure 2A**). Interestingly, these 5 genes were involved in immune modulation and have been previously identified in other studies as predictive biomarkers associated with responders to ICIs. For example, *CCL4* has been previously identified as a biomarker in the 12-chemokine signature (13,14), as well as other gene signatures predictive of neoadjuvant ipilimumab response (27). *IDO1* has been identified as a key marker in the IFN-γ signature (12) and gene signature predictive of response to ICIs in lung cancer (28). *CD96* is a marker that estimates CD8+ T cell infiltration (29,30). *CD96* and *TIGIT* along with the co-stimulatory receptor *CD226* form a pathway that affects the immune response in an analogous way to the CD28/CTLA-4 pathway (31). *CLEC12A* (32,33) and *GBP1P1* (34,35) were identified in immune-related gene expression signatures predictive of ICI responses.

**Figure 2.**
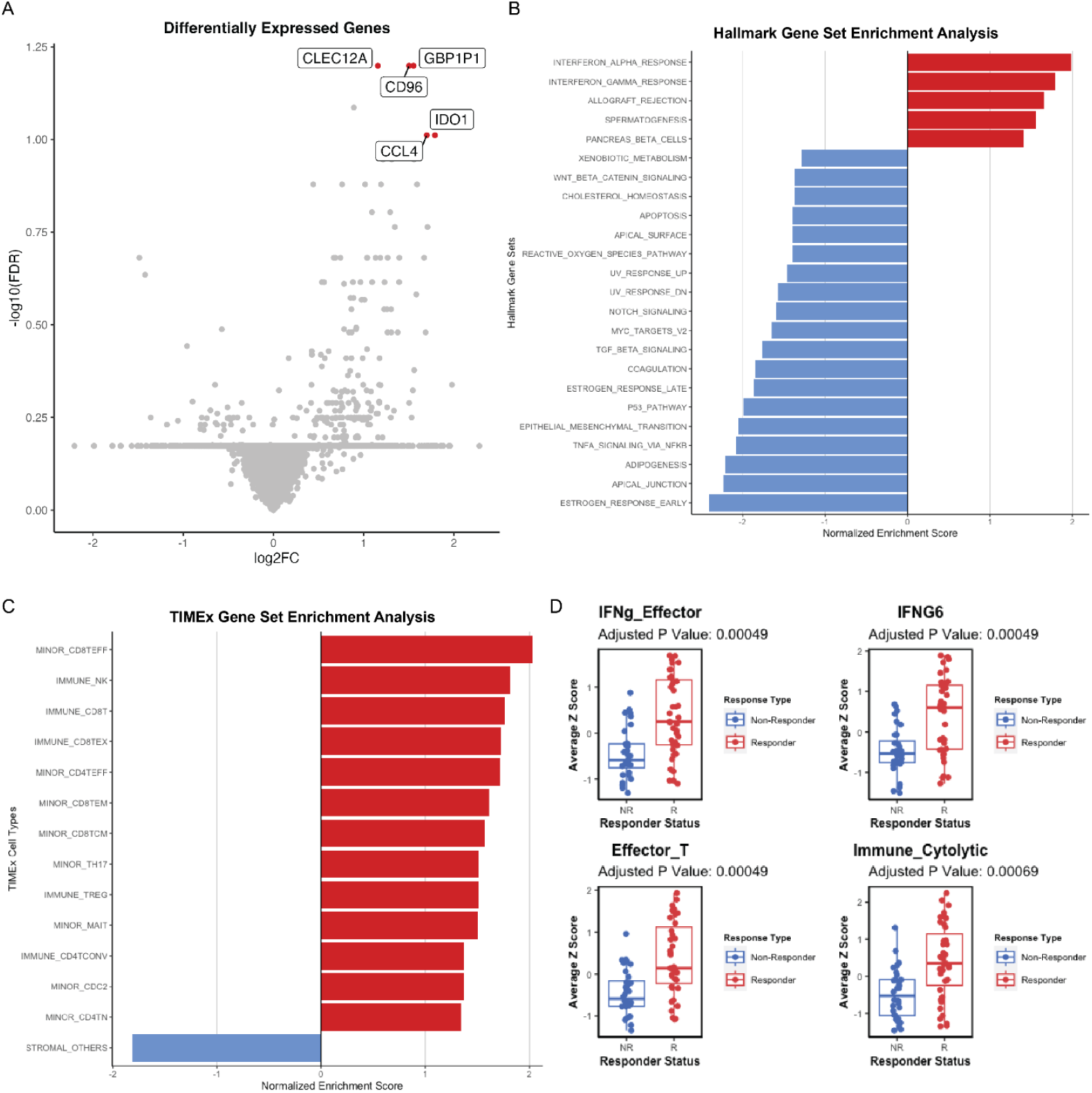
Immune-related gene expression associates with the response to ICIs. **(A)** Gene expression differences between the tumors that were responsive (right) and non-responsive (left) to ICI treatment. Significantly different genes after FDR correction are colored and labeled. **(B) and (C)** Gene set enrichment analysis comparing responders vs. non-responders using the Hallmark gene set and TIMEx cell types. FDR <0.1 was used as a cutoff. **(D)** Mann Whitney comparison of responders and non-responders for signatures reaching the 0.05 FDR threshold.

Next, we asked what gene sets and pathways were enriched or depleted in responders to ICIs. We performed GSEA using the MSigDB Hallmark gene sets on the RNA-seq and found that several immune-related gene sets were significantly enriched in responders (**Figure 2B**), for example, IFN-α response (NES = 1.98, FDR < 0.001), IFN-γ response (NES = 1.79, FDR < 0.001), and allograft rejection (NES = 1.65, FDR = 0.002). The other two gene sets enriched in responders were spermatogenesis (NES = 1.56, FDR = 0.005) and the pancreas beta cell gene sets (NES = 1.40, FDR = 0.036). In contrast, many cell-intrinsic gene sets were enriched in ICI non-responders as shown in **Figure 2B**. The GSEA results identified in this cohort are similar to previously published studies (23).

We next hypothesized that tumor-infiltrating immune cells could associate with responses to ICIs. To test this hypothesis, we performed cell-type deconvolution of the bulk RNA-seq using CIBERSORT. From CIBERSORT results, we observed that responders had significantly (p-value <0.05) higher abundances of CD8+ T-cells, activated CD4+ memory T-cells, activated NK cells, and M1 macrophages relative to non-responders, who were shown to have a significantly higher amount of resting mast cells **(Suppl. Figure 1)**. Similarly, when we performed GSEA using TIMEx gene sets, we observed that 13 CD4+, CD8+, and NK-related cell types were enriched in responders (FDR < 0.1), whereas the stromal cell type was enriched in non-responders (**Figure 2C**). This suggests that the tumor microenvironment of responders had an “immune-inflamed” phenotype, whereas non-responders had either “immune-excluded” or “immune-desert” TME phenotypes.

**Supplementary Figure 1.**
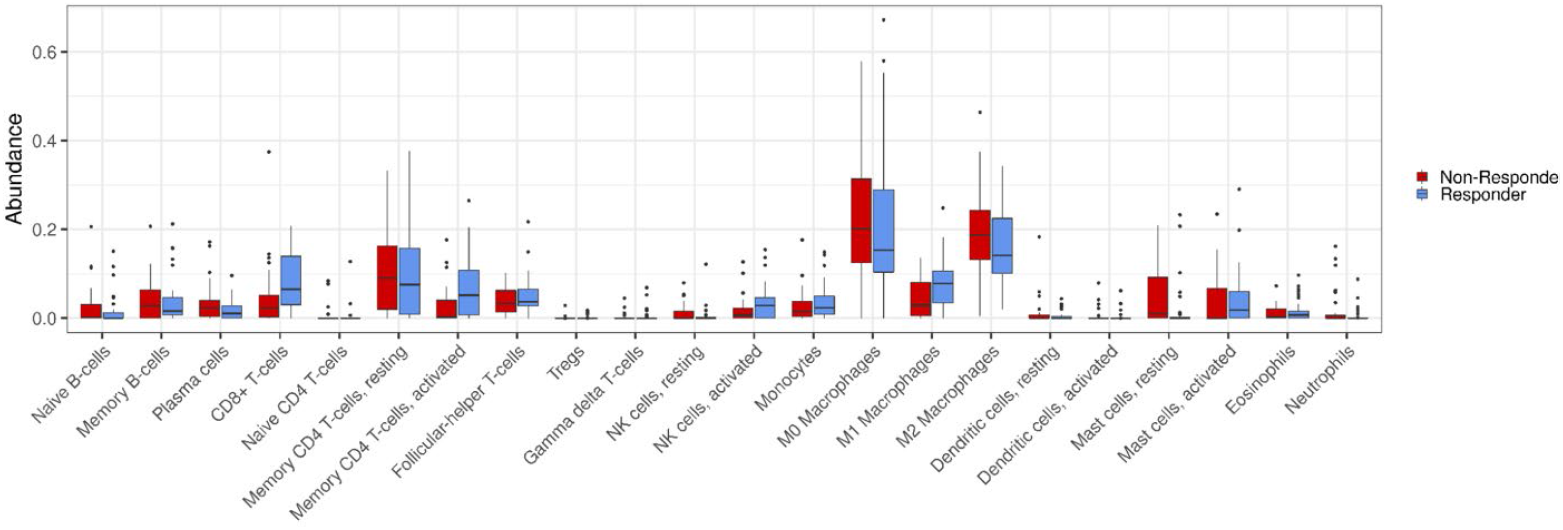
Association of CIBERSORT cell types with the response to ICIs.

**Supplementary Figure 2.**
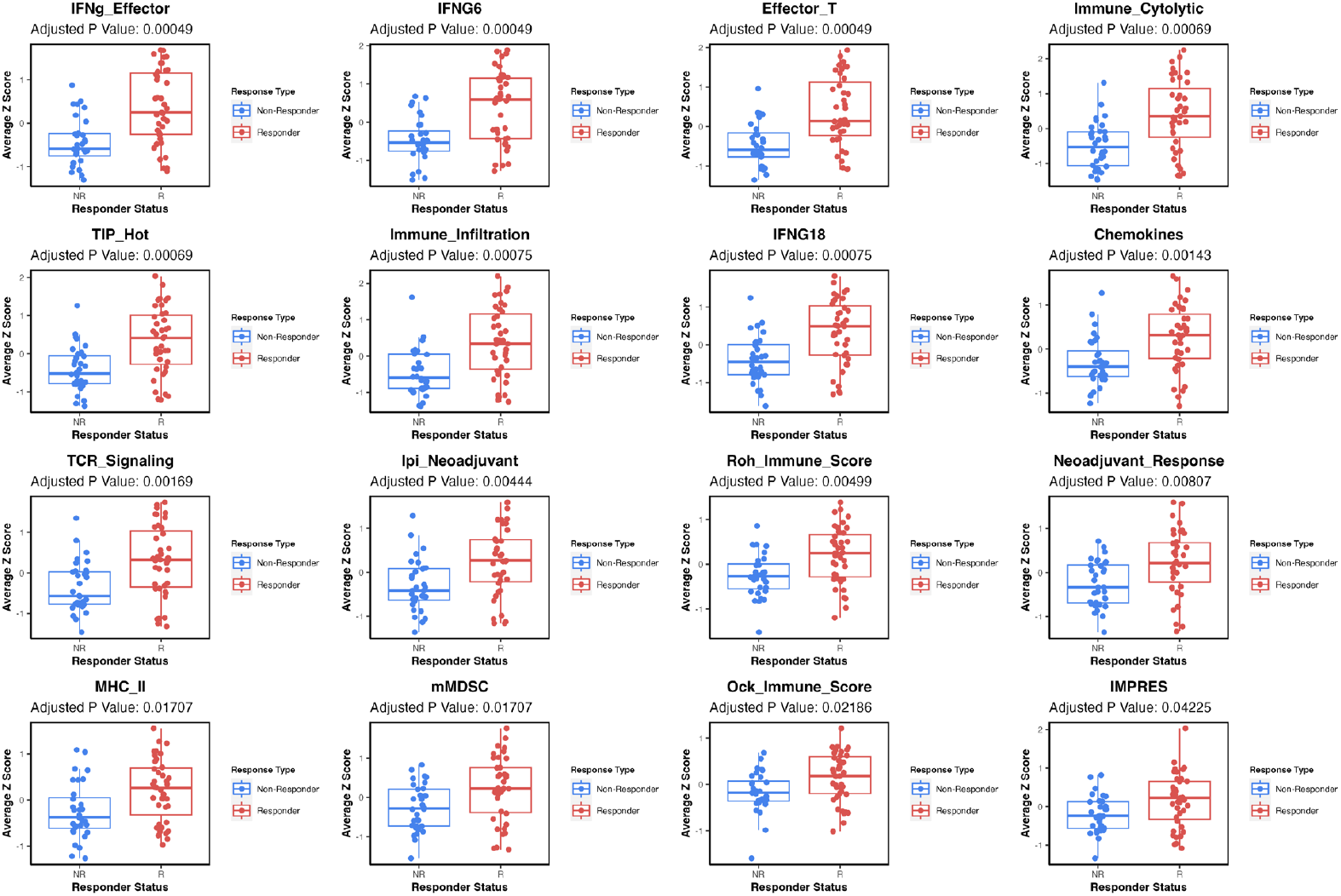
16 gene signatures where high Z-scores are associated with ICIs responsiveness in this cohort (FDR <0.05).

**Supplementary Figure 3.**
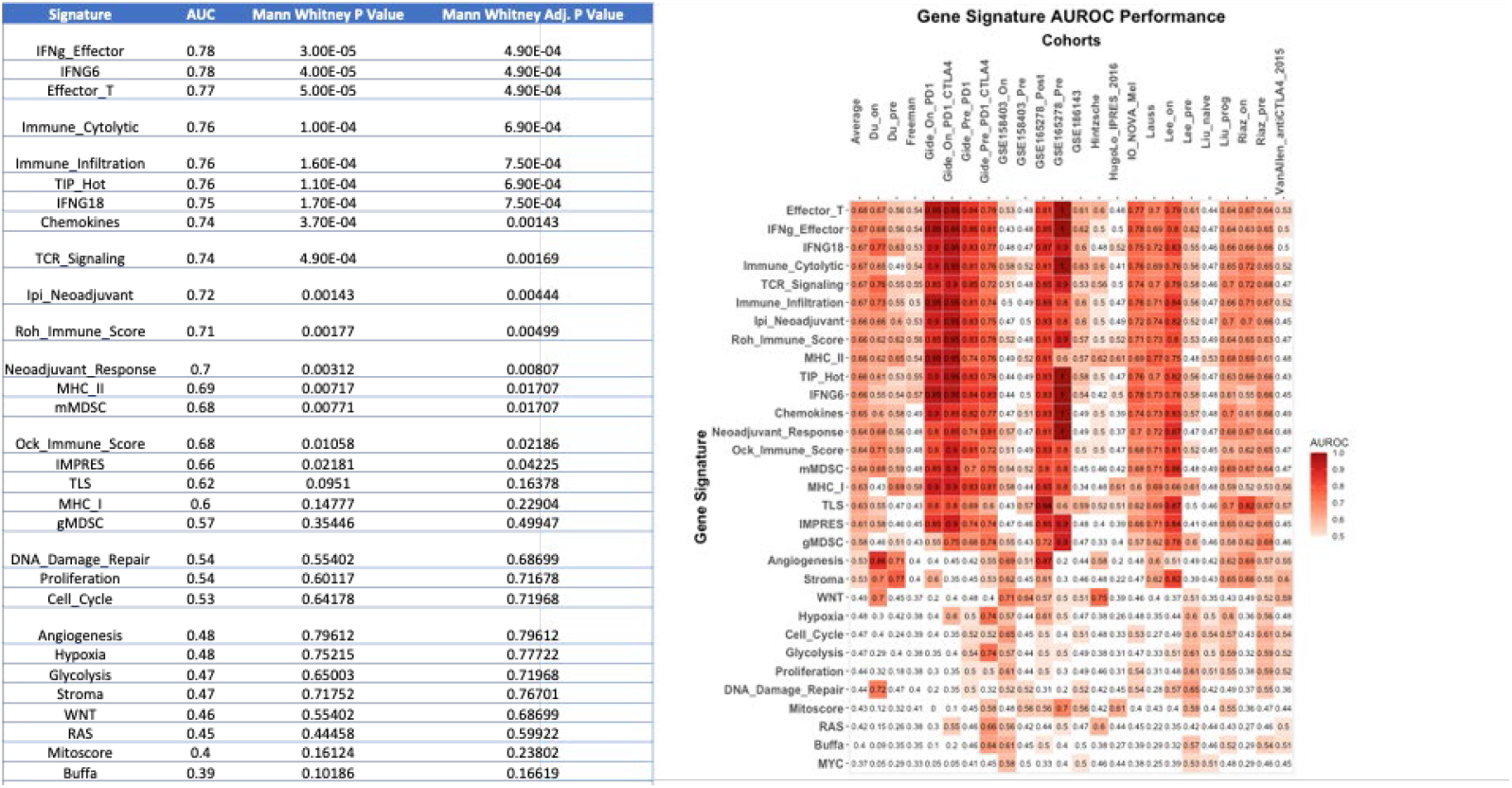
Predictive values (AUROC) of the 16 gene signatures in the ORIEN cohort (IO_NOVA_Mel) and 22 other melanoma cohorts.

To further delineate the immune phenotypes of responders vs. non-responders of ICIs, we used previously published gene signatures. We collected 30 gene expression signatures from the literature that have been implicated to be predictive of ICIs (23). By performing a Z-score for each signature and associating them with responders vs. non-responders, we identified 16 gene signatures (**Supplementary Figure 2**) where high Z-scores are associated with ICIs responsiveness in this cohort (FDR <0.05), and the top 4 gene signatures were illustrated in **Figure 2D**. These 16 gene signatures were related to immune activation and inflammation signatures (**Supplementary Figure 2**) (23).

We next used our recently developed IOSig portal (23) to evaluate the predictive values of these 16 gene signatures in our ORIEN cohort (IO_NOVA_Mel), as well as 22 other melanoma cohorts treated with ICIs. We used AUROC to assess the predictive value of these signatures. For the 16 gene signatures, the AUROC ranged from 0.78 to 0.66 in the IO_NOVA_Mel cohort (**Supplementary Figure 3**). On average, the AUROCs for these 16 gene signatures ranged from 0.61 to 0.68 in the separate 22 melanoma cohorts (**Supplementary Figure 3**).

### The melanoma tumor microbiome and its association with response to ICIs

Exogenous taxa were identified in the tumor RNA-seq, including bacteria, fungi, and viruses. A total of 54 phyla were observed, with *Firmicutes* being the most abundant phylum, followed by *Uroviricota* (**Figure 3A)**. Within the tumors responsive to immunotherapy, we found a significant enrichment of several microbes, including *Fusobacterium nucleatum, Porphyromonas asaccharolytica, Nocardia mangyaensis*, and *Mollivirus sibericum*. Comparatively, the cohort of non-responsive tumors was found to have significant intratumoral enrichment of fungi and the bacteria *Delftia lacustris, Enterobacter hormaechei, Pseudomonas fluorescens*, and *Moraxella osloensis* (**Figure 3B**). We observed no significant differences between alpha diversity metrics of responders and non-responders (Welch 2 sample t-tests p-value >0.4) (**Figure 3C**). We found that the random forest classifiers based on microbe diversity measures with 5 rounds of 5-fold cross-validation performed poorly relative to our other microbe-based classifers (**Figure 3D**).

**Figure 3.**
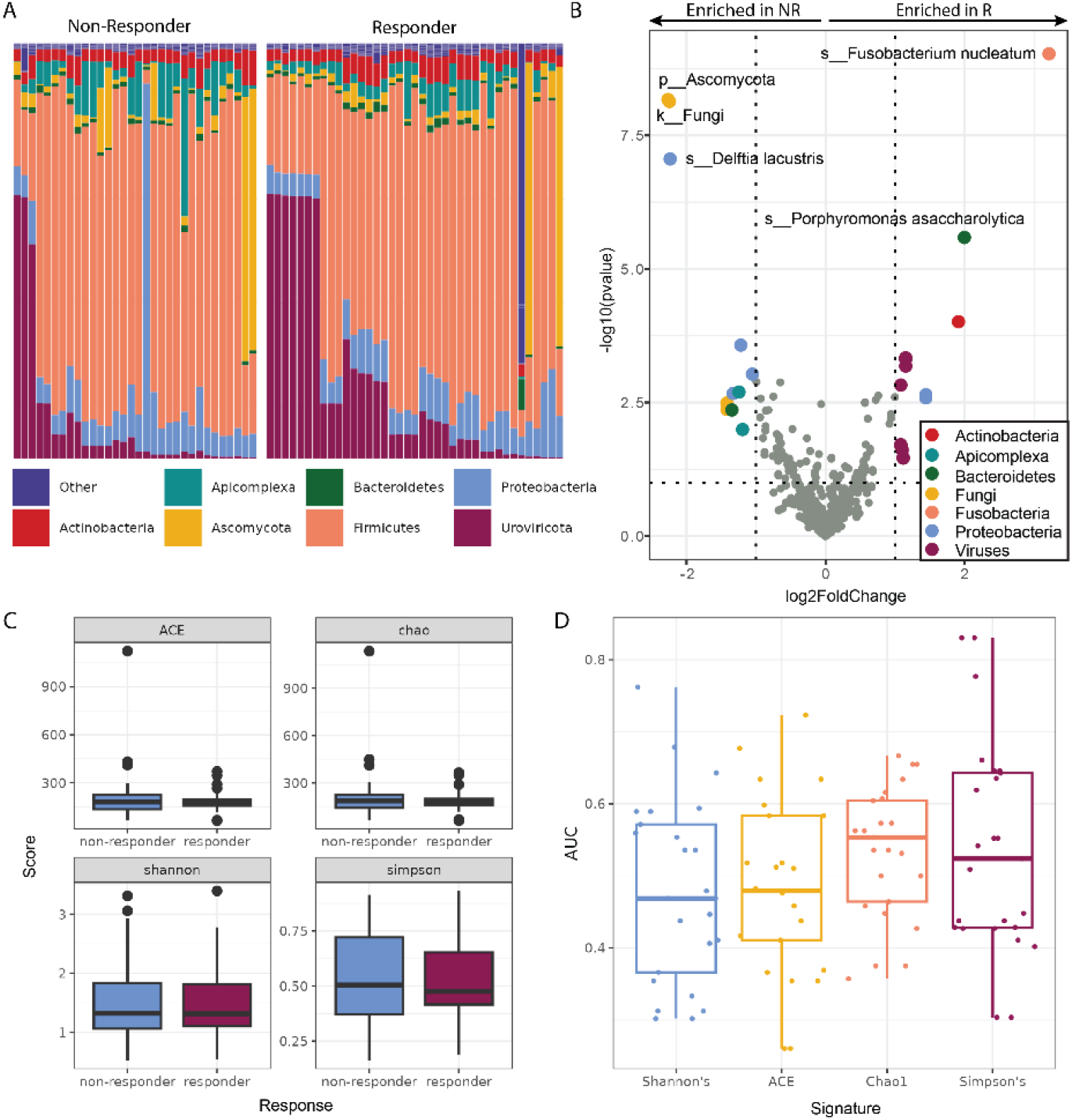
Melanoma tumors that respond to ICIs have a distinct tumor microbiome. **(A)** The relative abundances of the tumor microbiome at the phylum level showed wide intersample variation in the abundances of fungi (*Ascomycota* (yellow)) and viruses (*Uroviricota* (maroon)), without gross differences between non-responders (NR) and responders (R). **(B)** Differential abundance analysis of taxa found within tumor RNA-seq data. Colored points represent significantly (p-value <0.05) enriched taxa with a high (>1.00) fold difference in abundance between responders and non-responders to ICIs. **(C)** The diversity of the tumor microbiome between responders and non-responders shows no significant differences. **(D)** The diversity of the microbiome is a poor predictor of outcomes.

### Correlation of tumor RNA-seq (GSEA) with microbes

We next asked whether microbe abundance in the tumor could be associated with tumor intrinsic pathways or the composition of cell types in the tumor immune microenvironment. We focused on the 15 microbes identified to be differentially expressed in relation to immunotherapy response in melanoma. For the 15 microbes, 7 and 8 were associated with responders and non-responders of immunotherapy, respectively (**Figure 4A**). To investigate the intrinsic pathways that correlated with the microbes, we performed ssGSEA on melanoma patients using MSigDB

**Figure 4.**
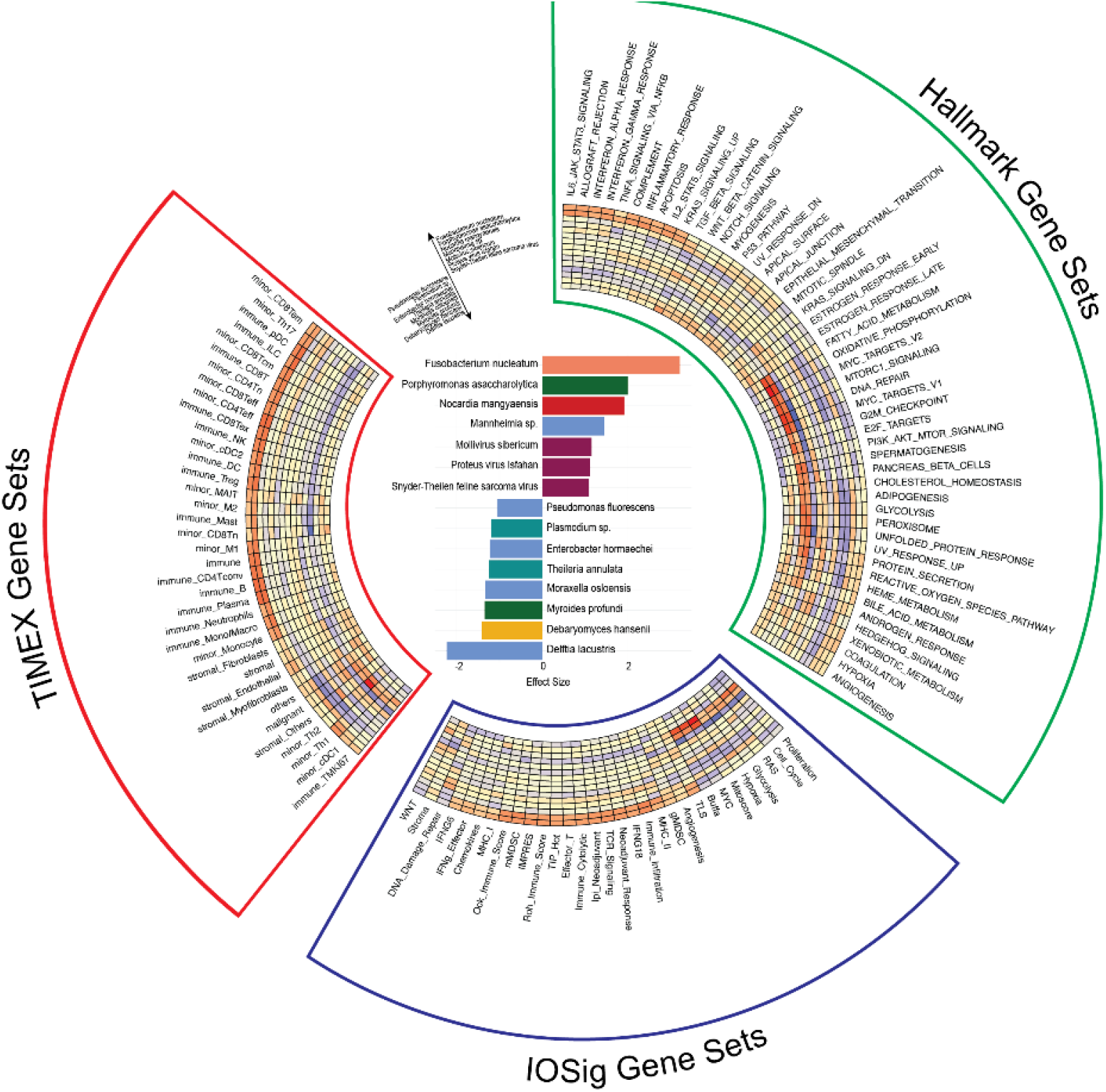
Association of microbes and gene signatures. **(A)** Effect size plot showing the top 15 most significantly enriched species. **(B), (C), and (D)** Spearman correlation coefficients between the significantly enriched species and the most significantly correlated signatures, and other gene sets, shown in a heatmap.

Hallmark gene sets. Interestingly, we observed that the two microbes highly abundant in responders, *Fusobacterium nucleatum* and *Porphyromonas asaccharolytica*, were correlated with inflammation and immune-related gene sets and pathways (**Figure 4B**). Conversely, we observed that the microbes that were highly abundant in non-responders, *Theileria annulata* and *Moraxella osloensis*, were correlated with intrinsic gene regulation (e.g., MYC target gene sets, E2F target genes), DNA damage repairs, intrinsic cell signaling pathways (e.g., MTORC1 signaling, PI3K-AKT-MTOR signaling), and metabolisms (e.g., fatty acid metabolism, glycolysis) (**Figure 4B**). These results are consistent with our previous findings, where we observed the same Hallmark gene sets and pathways enriched in responders vs. non-responders across 5 melanoma cohorts of immunotherapy-treated patients with pre-treatment and on-treatment tumor biopsies (23).

To further dissect the association of microbe abundance and the composition of cell types in the context of immunotherapy responses in melanoma, we performed cell type deconvolution using the bulk RNA-seq with TIMEx. We found that the two microbes, *Fusobacterium nucleatum* and *Porphyromonas asaccharolytica*, were highly correlated with the enrichment of tumor-infiltrated immune cell types, including CD8+ T cells, which are known predictors of immunotherapy response (**Figure 4C**). In contrast, the lack of tumor-infiltrated immune cell types was correlated with microbes associated with non-responders. In particular, we observed that malignant and stromal cell types were enriched in association with the 2 tumor microbes noted in non-responders, *Theileria annulata* and *Moraxella osloensis* (**Figure 4C**). The tumor immune cell composition corroborated our previous findings (23).

Next, we asked whether the microbe abundance was associated with any gene signatures predictive of immunotherapy responses. To investigate this question, we utilized 31 previously published gene signatures that have been indicated to be associated with immunotherapy responses (23). We correlated microbe abundance with these signatures, and found that gene signatures associated with inflammation or immune activation were highly associated with microbes abundant in responders (**Figure 4D**). On the other hand, gene signatures associated with immune-suppressive or intrinsic signaling were highly associated with microbes abundant in non-responders (**Figure 4D**). These results suggest that microbe abundance could provide a different dimension in understanding the tumor immune microenvironment in predicting immunotherapy responsiveness in melanoma.

### Prediction of response using tumor gene expression and microbe abundance

We further hypothesized that combining microbe abundance features with gene expression signatures could improve response prediction of melanoma to immunotherapy. To test this hypothesis, we developed an ensemble learning random forest classifier using microbe abundance and gene signatures identified to be associated with immunotherapy responses in melanoma. We first developed the random forest classifier based on microbe abundance with 15 input features (microbe) and performed 5 rounds of 5-fold cross-validation on the melanoma cohort (**Figure 5**). The average AUROC for the microbe classifier was 0.651. We also constructed a random forest classifier based on 31 gene signatures (GeneSig_Z_score) or the 16 immune-activated gene signatures (Imm_Act_Z_score), and the AUROC values for GeneSig or Imm_Act classifiers were 0.72 and 0.744, respectively (**Figure 5**). Notably, when we combined the microbe abundance and gene signatures to develop the random forest classifier, the ensemble learning random forest classifiers for gene signatures plus microbe (GS_Z_microbe) and immune-activated gene signatures plus microbe (Imm_Act_Z_microbe) achieved 0.772 and 0.805, respectively (**Figure 5**). This suggests that microbe abundance features provide a distinct layer of information in predicting response to immunotherapy and, when combined with gene expression signatures, can improve the prediction of response to immunotherapy in melanoma.

**Figure 5.**
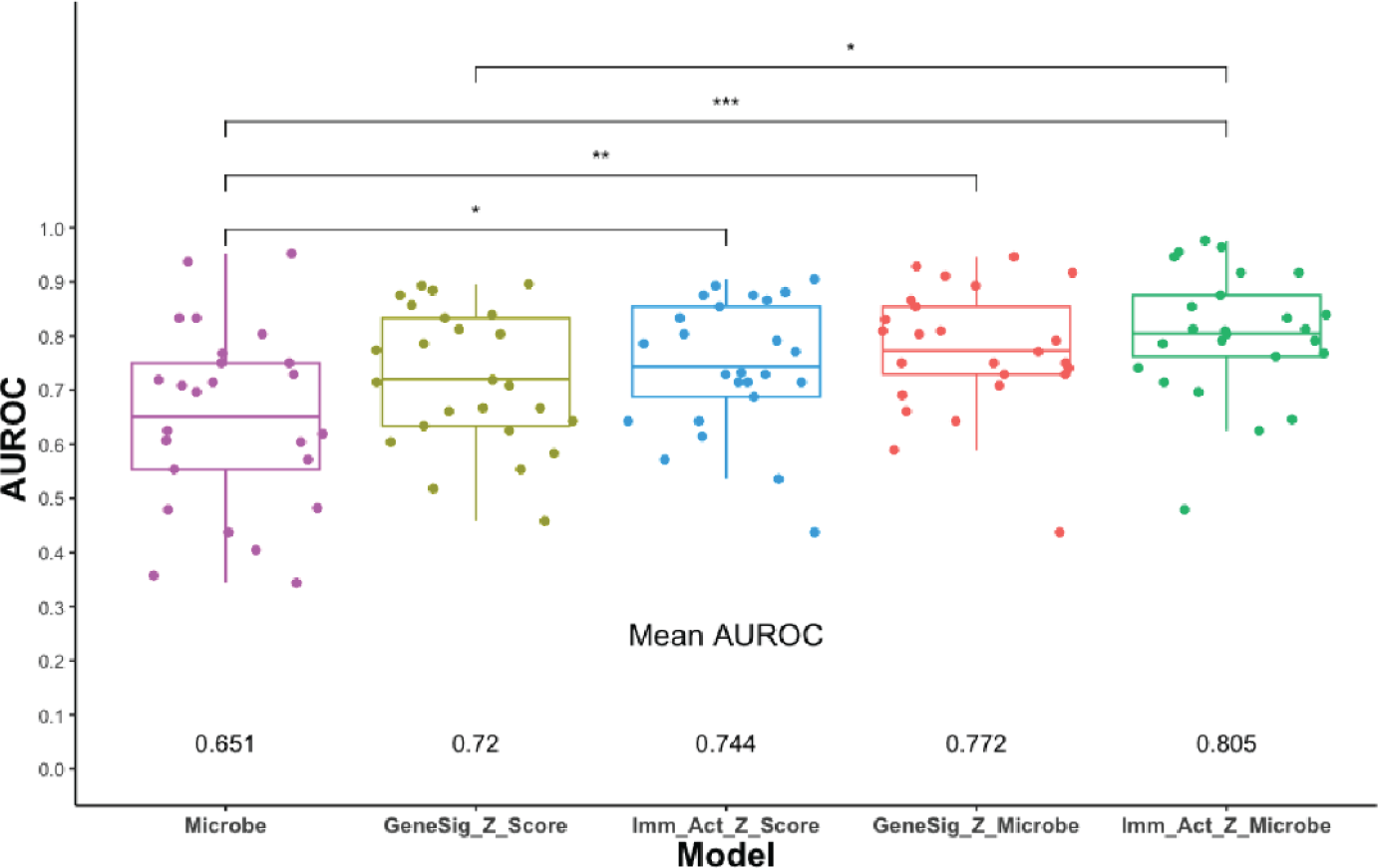
Prediction of response using gene expression and microbes. Mann Whitney comparisons of the mean AUROCs from random forest model comparisons

## DISCUSSION

We utilized tumor RNA-seq from melanoma patients to explore the tumor microbiome’s influence on clinical outcomes, specifically in response to ICIs. We observed microbes in all samples, and showed that tumors that responded to ICIs had significantly different taxa present from those that didn’t respond to treatment. Consistent with previous findings, gene expression seems to be predictive of response to ICIs. In addition, we showed that microbes are also predictive of response to ICIs, particularly when combined with gene expression, suggesting that the inclusion of microbes in these models enhances predictive ability.

A correlation between the gut microbiome and response to ICIs has been consistently indicated in previous research (36–38). Altering the gut microbiome via responder-derived fecal microbiota transplantation has been shown to induce a clinical response to anti-PD-1 treatment in melanoma patients (39,40). However, many of the efforts in this area have focused solely on the gut microbiome. Therefore, we assessed the tumor microbiome to further explore the impact of microbes on clinical outcomes in body sites beyond the gut.

We observed the presence of microbiota in all 71 tumor samples, as is consistent with previous findings regarding the tumor microbiome (41,42). Our study explicitly exhibits the microbial characteristics of tumors in patients with metastatic melanoma. Previous research has shown that the tumor microbiome in this specific subset of cancer is predictive of response to treatment, but these findings have been limited in scope due to samples having been collected before the use of modern ICIs as a standard treatment regimen for metastatic melanoma (20). We showed distinct, significantly enriched taxa, including fungi, at baseline for patients treated with contemporary ICI-based treatment plans.

The mechanisms by which tumor microbes affect response to ICIs may relate to interactions with the immune system or several other established mechanisms (43).The World Health Organization (WHO) has officially recognized a causal association between 11 microbes and cancer (44). However, in recent years, the number of likely carcinogenic microbes and more loosely related “complicit” microbes has increased dramatically. These have been shown to interact with the host via diverse mechanisms. For example, in colon cancer, *Bacteroides fragilis* biofilms on colon polyps have been found to secrete a toxin that directly damages DNA (45,46), as have some *Escherichia coli* (47). In another mechanism, *Helicobacter pylori* secrete a series of molecules eliciting an inflammatory cascade shown to drive tumorigenesis in gastric adenocarcinoma and mucosa-associated lymphoma (48,49). The fungal genus *Malassezia* caused pancreatic ductal adenocarcinoma growth through activation of the C3 complement pathway (50). Several microbes enriched in responders have strong precedence for interacting with the human immune system. *Fusobacterium nucleatum*, which correlated most strongly with responders, has been shown to increase tumor growth rates in colorectal cancer (51), as it produces a pro-inflammatory microenvironment favorable to tumor growth (52,53). On the other hand, *Porphorymonas* has not been associated with the tumor microbiome or response to ICIs although it is an established pathogen that has been linked to colorectal cancer (54). In our study, it is associated with the same immune expression pathways as *Fusobacterium nucleatum*, suggesting it acts through a similar mechanism. The diversity of mechanisms and taxa suggests that additional mechanisms are likely. Furthermore, recent studies have identified bacteria-derived human leucocyte antigen (HLA)-bound peptides in melanoma presented by tumor cells could elicit immune reactivity. This intratumoral bacteria peptide repertoire could be further explored to understand the mechanism by which bacteria modulate the immune system and responses to therapy (55). The demonstration of the utility of high-throughput sequencing to explore these correlations warrants a broader search.

Efforts have been made to identify predictors of response and resistance to ICIs. As previously discussed, expression signatures have been established as predictors of ICI response in metastatic melanoma (9,12,14,15,23,56). One such study assessing the model combining IFNγ and TMB found that it was predicitve of response but not resistance (56). Another such study developed a multi-omic-based classifier that successfully predicted response, but was also unable to predict resistance (20). We showed significantly enriched taxa in both response groups. We also showed that microbes alone are predictive of response/resistance to immunotherapy and, when combined with gene expression, enhance the model’s predictive ability. Further studies are warranted to combine tumor microbiome abundance with other clinical and “omics” (e.g., genomics and pathomics) for developing an accurate classifier for predicting immunotherapy responses in melanoma. Our findings also warrant further research to evaluate whether these correlations are causally associated with outcomes and their effect on the tumor immune microenvironment and immune cell infiltration.

In conclusion, we found that the tumor microbiome in patients with metastatic melanoma was significantly different in those that responded (>24 months survival) to treatment with ICIs from those who didn’t respond. Furthermore, the microbial communities had the ability to predict response when incorporated into machine learning models. The tumor microbiome further enhanced models to predict response when combined with gene expression data. Future research has the potential to support therapeutic strategies to modify the tumor microbiome to improve ICI treatment outcomes.

## ACKNOWLEDGMENTS

The authors acknowledge the support and resources of the Ohio Supercomputer Center (PAS1695). We would like to thank Angela Dahlberg, Editor, Division of Medical Oncology at The Ohio State University Comprehensive Cancer Center, for editing and proofreading the manuscript.

